# DynamoSort: Using machine learning approaches for the automatic classification of seizure dynamotypes

**DOI:** 10.1101/2025.02.12.637999

**Authors:** Josh Wooley, Ashley Zachery-Savella, Michelle Le, Sally Scofield, Kishore Jay, Josh Mosse-Robinson, Peter J. West, Karen S. Wilcox, Daria Nesterovich Anderson

## Abstract

**Objective:** Epilepsy is characterised by unprovoked and recurring seizures, which can be electrically measured using electroencephalograms (EEG). To better understand the underlying mechanisms of seizures, researchers are exploring their temporal dynamics through the lens of dynamical systems modelling. Seizure initiation and termination patterns of spiking amplitude and frequency can be sorted into “dynamotypes”, which may be able to serve as biomarkers for intervention. However, manual classification of these dynamotypes requires trained raters and is prone to variability. To address this, we developed DynamoSort, a machine-learning algorithm for automatic seizure onset and offset classification.

**Methods:** We used approximately 2100 seizures from an intra-amygdala kainic acid (IAK) mouse model of mesial temporal lobe epilepsy, categorized by five trained raters. MATLAB’s classification learner application was used to create an ensemble model to score and label dynamotypes of individual seizures based on spiking and frequency features.

**Results:** Dynamotype classification of real EEG data lacks a definitive ground truth, with mean inter-rater agreement at 73.4% for onset and 64.2% for offset types. Despite this, DynamoSort achieved a mean area under the curve (AUC) of 0.81 for onset and a mean AUC of 0.75 for offset types. Machine-human agreement was not significantly different from human-to-human agreement. To address the lack of ground truth in ratings, DynamoSort assigns probabilistic scores (-20 to 20), to indicate similarity to each seizure dynamotype based on spiking features, allowing for a characterization of seizure dynamics on a spectrum rather than the traditional qualitative taxonomy.

**Significance:** Automating the classification of dynamotypes is a critical step for their inclusion as a biomarker in clinical and research applications. DynamoSort is a straightforward, open-access tool that uses automatic labelling and probabilistic scoring to quantify subtle changes in seizure onset and offset dynamics.

**Key Points:**

- Dynamotypes are a promising seizure categorization system, but is prone to interrater variability and lacks a ground truth.
- Machine learning can be used to automatically classify seizure onsets and offsets into appropriate dynamotypes based on spike features.
- Agreement between DynamoSort and human raters rivals typical agreement rates in trained human raters.
- DynamoSort uses probabilistic scoring to quantify subtle changes in seizure onset and offset, allowing for a quantitative characterisation.

## Introduction

Epilepsy is a neurological condition characterized by recurring seizures, affecting roughly 65 million people globally. An estimated one-third of those cases are treatment resistant (Moshé et al. 2015). Electroencephalography (EEG) is an essential tool for diagnosis and treatment of epilepsy (Noachtar & Rémi 2009). EEG also plays a role in biomarker discovery and subclassification of seizures based on their electrographic features. Categorizing seizure onset patterns, such as initial slow waves, low-voltage fast activity, or the presence of a DC-shift, can offer insight into mechanisms of seizure generation (Bragin et al. 2007, Elahian et al. 2018, Staba et al. 2014).

Recent approaches have employed dynamical systems modelling and bifurcation theory to define and classify seizure properties (Jirsa et al. 2014, Saggio et al. 2020, Saggio et al. 2017). A dynamotype is a description of the shape and evolution of the onset and offset— two distinct bifurcations—of a seizure based on stereotyped changes in spike amplitude or frequency. Although dynamotypes are a promising classification system, manually rating seizures is prone to interrater variability and can be time intensive. We developed DynamoSort—a freely available machine learning algorithm designed to automatically and accurately classify seizure onset and offset zones by bifurcation type.

While the original dynamotype taxonomy was developed for clinical utility, it has also been applied to rodent and *in vitro* seizure-like-events (Crisp, Cheung, et al. 2020, Crisp, Parent, et al. 2020). Rodent models are widely used in mechanistic epilepsy research and their seizures reliably recapitulate EEG properties of humans (Milior et al. 2023, Rusina et al. 2021), including specific dynamic properties such as high-frequency oscillations and aperiodic signal (Maheshwari 2020). Since dynamotype classification is based solely on spiking amplitude and frequency patterns, a machine learning model may be trained using EEG data agnostic of source, whether from simulation, rodent, or human. In this manuscript, we detail the development, training, and testing of DynamoSort, as well as its approach to classifying ambiguous dynamotypes in comparison to human raters.

## Methods

### Dynamotypes defined

According to the existing taxonomy (Saggio et al. 2020), eight bifurcation types can be defined: four at onset and four at offset (Figure 1). Of these eight types, one onset type and one offset type require a measurable baseline shift in the EEG, i.e. direct current shift (DC shift). If EEG data underwent DC shift filtering during recording, as in the datasets used in this analysis, only three onset and three offset bifurcation types can be meaningfully distinguished.

**Figure 1.**
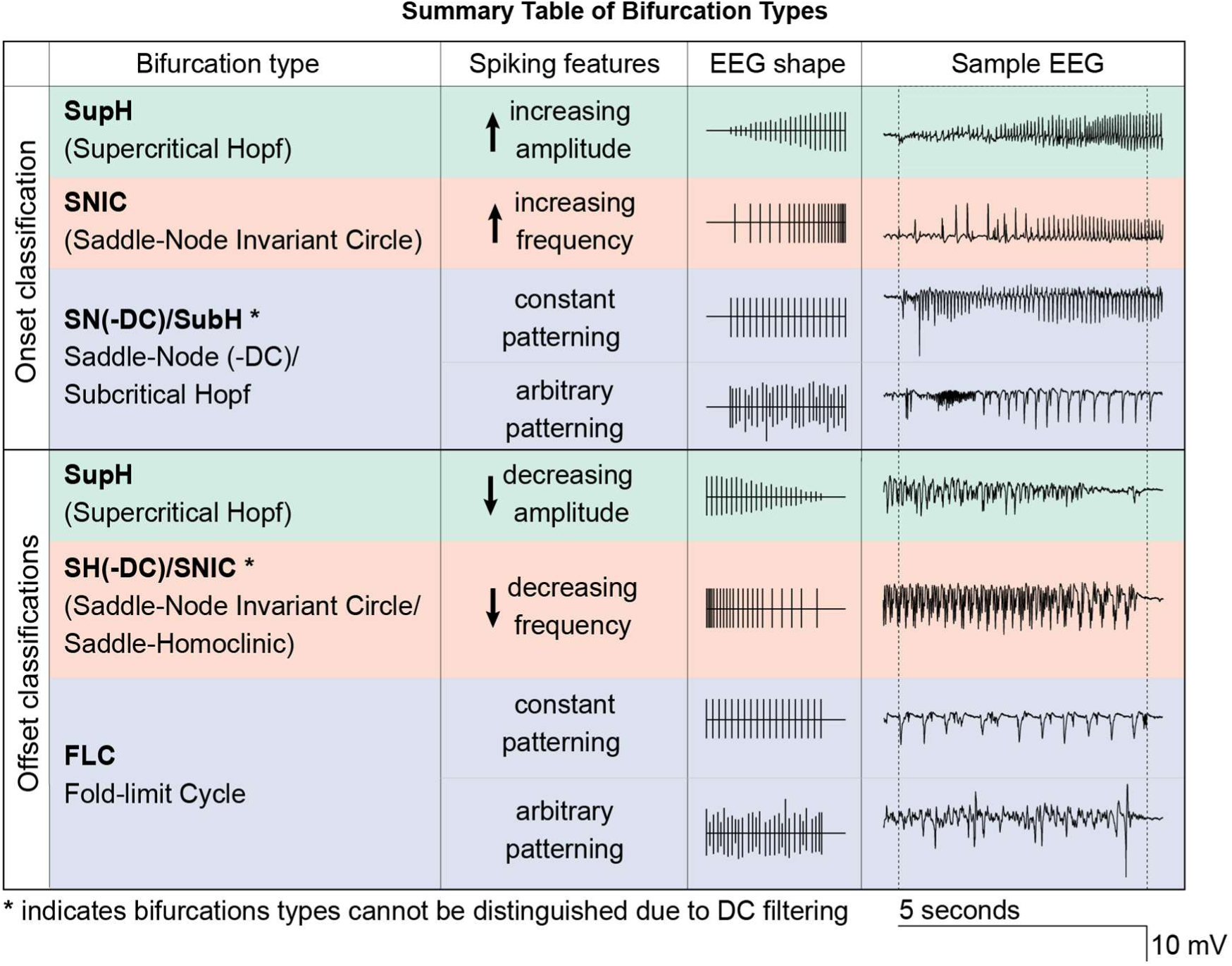
Summary table of bifurcation types. Full names and abbreviations of the bifurcation types used in the labelled data are shown, split into onset or offset categories. Characteristic spike features for each type are described, along with a representation of ideal shape and an exemplary EEG sample from the mouse seizures used in the study. Due to the filtering of the DC offset in the datasets used, both onset and offset have types that cannot be distinguished from each other: SN(-DC) and SubH for onset cannot be distinguished, and SH(-DC) and SNIC for offset cannot be distinguished. Additionally, SN(-DC)/SubH and FLC onset and offset types may exhibit constant or arbitrary spike patterning, but both subtypes are grouped together in this analysis because neither demonstrates the stereotyped change of amplitude or frequency.

Onset Supercritical Hopf (SupH) is defined by a stable frequency and a proportional increase in amplitude. Onset Saddle-Node on an Invariant Circle (SNIC) is defined by an increase in spiking frequency with a stable amplitude. Saddle-Node (SN) and Subcritical Hopf (SubH) are defined by either arbitrary or constant amplitude and frequency changes that do not follow a regular pattern seen in SupH and SNIC bifurcation types. Because SN typically displays a DC shift, not distinguishable in our study, they are referred to together as SN(-DC)/SubH.

Offset SupH is defined by a stable frequency and a proportional decrease in amplitude, the reverse of onset SupH. Both offset SNIC and offset Saddle-Homoclinic (SH) bifurcations are defined by a decreasing frequency, but offset SH is additionally differentiated by a DC shift. These types are referred together as SH(-DC)/SNIC. Finally, offset Fold Limit Cycle (FLC) may show either constant or arbitrary patterning in amplitude and frequency, often with an abrupt termination of the seizure.

### Datasets for training, validation, and testing

We obtained three annotated intra-amygdala kainic acid (IAK) mice seizure EEG datasets from the Epilepsy Therapy Screening Program contract site at the University of Utah for the training, validation, and testing of our machine learning classifier (Supplemental Table 1). In this dataset, chronic, intracranial EEG was used to record seizures after mice received kainic acid microinjections into the basolateral amygdala and developed spontaneous, recurrent seizures as a model of mesial temporal lobe epilepsy (West et al. 2022). From West et al. 2022, we were provided 2112 seizures captured across a 90-day epileptogenesis cohort and 2 anti-epileptic drug studies cohorts. These datasets, collected under varying conditions, help ensure our model is robust to a larger range of features.

The primary training cohort for DynamoSort was from a 90-day epileptogenesis (“epileptogenesis”) study (Supplement Table 1). The other two cohorts, part of a drug-crossover study, were treated with phenobarbital (Drug 1) or valproic acid (Drug 2). Both anti-seizure medications (ASMs) were effective at reducing seizure frequency in the IAK mouse (West et al. 2022). Since people with epilepsy often take ASMs during EEG monitoring, and future dynamotype studies will likely involve those on medication, this data was included to ensure the algorithm remained agnostic of dataset when testing. While ASMs could influence the prevalence of certain dynamotypes, classification is based on spiking and spectral features of 5-second onset and offset periods. The process itself is unaffected by ASM presence.

### Rating onset and offset dynamotypes

The onset period is defined as a five second window beginning at the start of each seizure, and the offset period spans from five seconds prior to seizure termination until its cessation. Raters were trained to label onsets as “SupH”, “SNIC”, “SN(-DC)/SubH” or “N/A”, and offsets as “SupH”, “SH(-DC)/SNIC”, “FLC”, or “N/A”. The “N/A” category includes onsets and offsets that raters could not reasonably classify, often due to signal quality degradation by 60 Hz line noise. During manual classification, human raters relied solely on the visual features of the seizure at onset and offset, such as increasing amplitude or decreasing frequency, to determine dynamotypes. All raters followed the classification approach outlined in Table 1 of the Appendix in Saggio et al., 2020 to independently classify dynamotypes in a randomized, blinded fashion. Given that there is no objective “ground truth” for dynamotype classification of real EEG data, it was important to train multiple raters to establish which bifurcation type was best suited to describe the onset or offset. Raters A, B, and C labelled all seizures within the epileptogenesis and Drug 1 datasets. Independently, raters D and E labelled the onsets and offsets in the Drug 2 dataset, providing additional validation of the classifier’s performance against raters not included in the training set (Supplemental Table 1).

### Feature extraction

To replicate the raters’ classification approach and the spike detection method described by Saggio et al., 2020, we extracted frequency and amplitude features from each EEG onset and offset signal. All scripts were written in MATLAB (The MathWorks Inc. 2023) and utilised custom code. Each recorded seizure, sampled at 500 Hz, was z-score normalised prior to feature extraction. To extract spiking features for both positive and negative-going peaks, the instantaneous amplitude (the envelope) of the signal was calculated by taking the absolute value of the Hilbert transform of the EEG signal (Figure 2A). A “smoothed” envelope was also generated using the *movmean* function (window size = 51; smoothing over 100 ms) to account for bursting activity, where multiple spikes may be grouped as a single burst for frequency calculations (Saggio et al. 2020). Subsequently, the *findpeaks* (minimum peak prominence = 0.1 and 0.75) function in MATLAB was used to extract spike amplitude and their corresponding timings.

**Figure 2.**
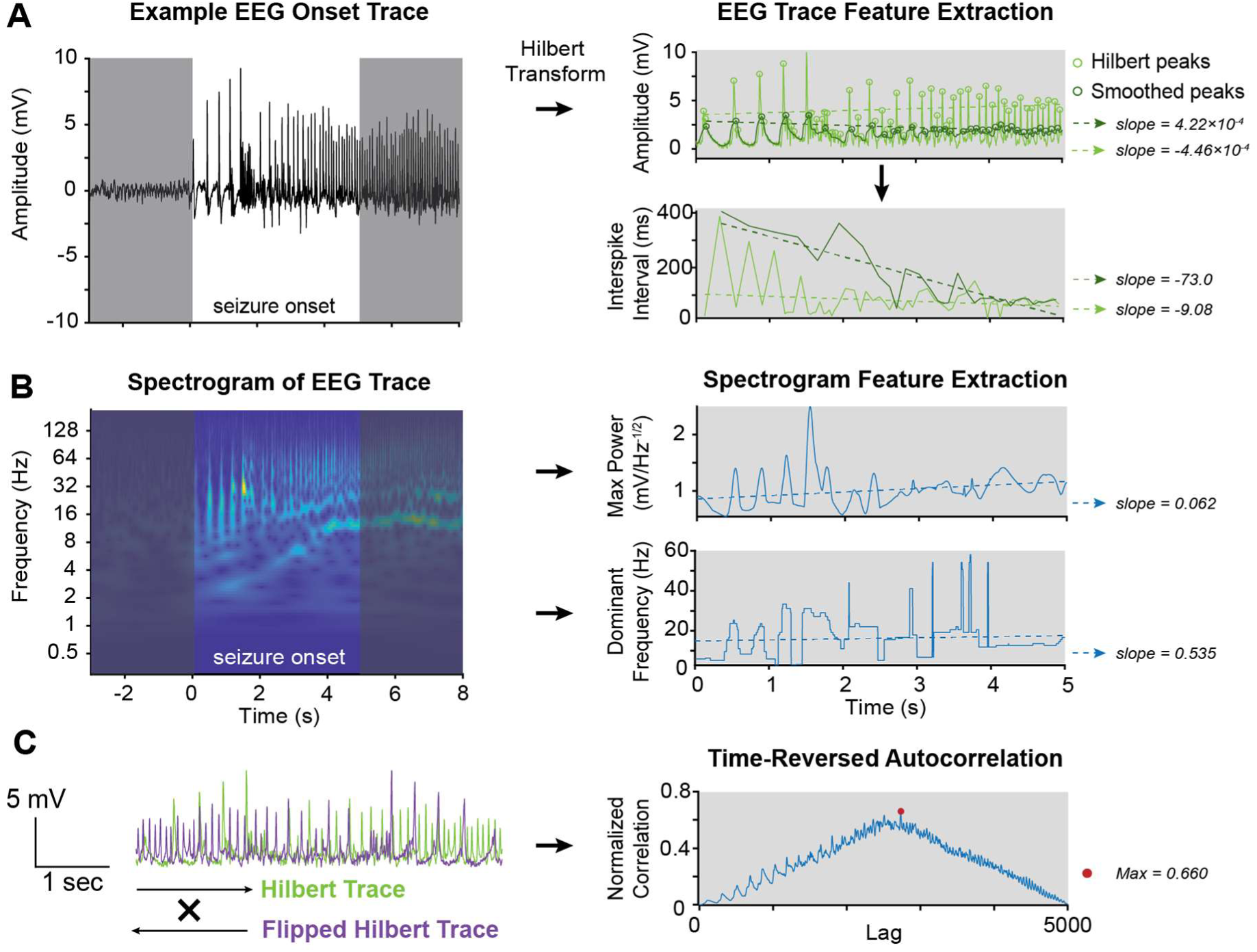
EEG Feature Extraction. **(A)** An example onset EEG trace from a mouse in the epileptogenesis dataset along with slope features extracted from the Hilbert envelope (both raw and smoothed). The raw and smoothed envelope was run through peak detection function, and linear fits were applied to the amplitude inter-spike interval over time to quantify the change in amplitude and frequency of the signal. **(B)** The continuous wavelet transform was visualised to demonstrate power across different frequencies of the EEG trace from 2A. The maximum power, and frequency at which the maximum power occurs, from each timepoint was plotted against time, and linear fits were generated as model features to quantify changes in amplitude and frequency. **(C)** The time-reversed autocorrelation of the Hilbert envelope was calculated, and the maximum value was extracted to capture the similarity of the signal to itself.

We applied linear fits to quantify changes in amplitude and interspike interval over onset and offset periods (Figure 2A), and their slopes were used as features to train and test the classifier model. Continuous wavelet transformations (*cwt* function) were calculated, and linear fits were used to quantify change in the rate at which the dominant frequency and the power at the dominant frequency changed during the 5second onset and offset period (Figure 2B). R^2^ values from all slope calculations were included as additional features used by the algorithm.

To better capture arbitrary spiking patterns, we performed a time-reversed autocorrelation on the Hilbert envelope and extracted the normalized maximum correlation (Figure 2C), where higher values indicated greater similarity between the beginning and end of the signal. In total, DynamoSort extracts 17 total features for dynamotype classification.

### Machine learning model selection

We used the MATLAB Statistics and Machine Learning Toolbox Classification Learner to test a variety of machine learning models to find the best performer for our training dataset. The criteria in building our model were that it be robust, easy to train, relatively fast, and be able to perform at least as well as we would expect a trained human rater to perform. Using the extracted features, we began by training all classifier types provided in the toolbox. Four classifiers were selected based on performance to move on to more rigorous training and testing: *k*-nearest neighbour, naïve bayes, ensemble classifier, and a neural network. For each, the receiver operating characteristic curve (ROC curve) and the area under the curve (AUC) were calculated to define performance. After rounds of training and 10-fold cross-validation, the best classifier was chosen based on highest AUC and accuracy, standard measurements of model performance (Jin Huang & Ling 2005).

### Unbalanced bifurcation labels and disagreement

To address anticipated disagreement between raters on bifurcation classification in the training set, seizures were only included if at least two raters agreed on its classification. Seizures were excluded in rare cases with complete rater disagreement. To balance the training data, seizures with full rater agreement were also added to a separate list that was used for oversampling. Full agreement cases were considered more representative of each bifurcation type and served as better training examples. Oversampling was performed by randomly selecting from this list until the class sizes were even.

### Training, validating, and testing the classifier

Once the classifier type was chosen, a model was trained and validated using 10-fold cross-validation for each bifurcation type individually and then combined, as shown in Figure 3B. However, models for onset and offset classification were kept separate–the most consistent and accurate approach, allowing hyperparameters to be flexible based on bifurcation type. Using bifurcation type-specific models allows the generation of a “likeness” score for each type, the maximum of which determines the bifurcation label. This allows the model to give a discrete label answer as well as a score on a continuum to capture when onsets and offsets may have features of multiple bifurcation types.

**Figure 3.**
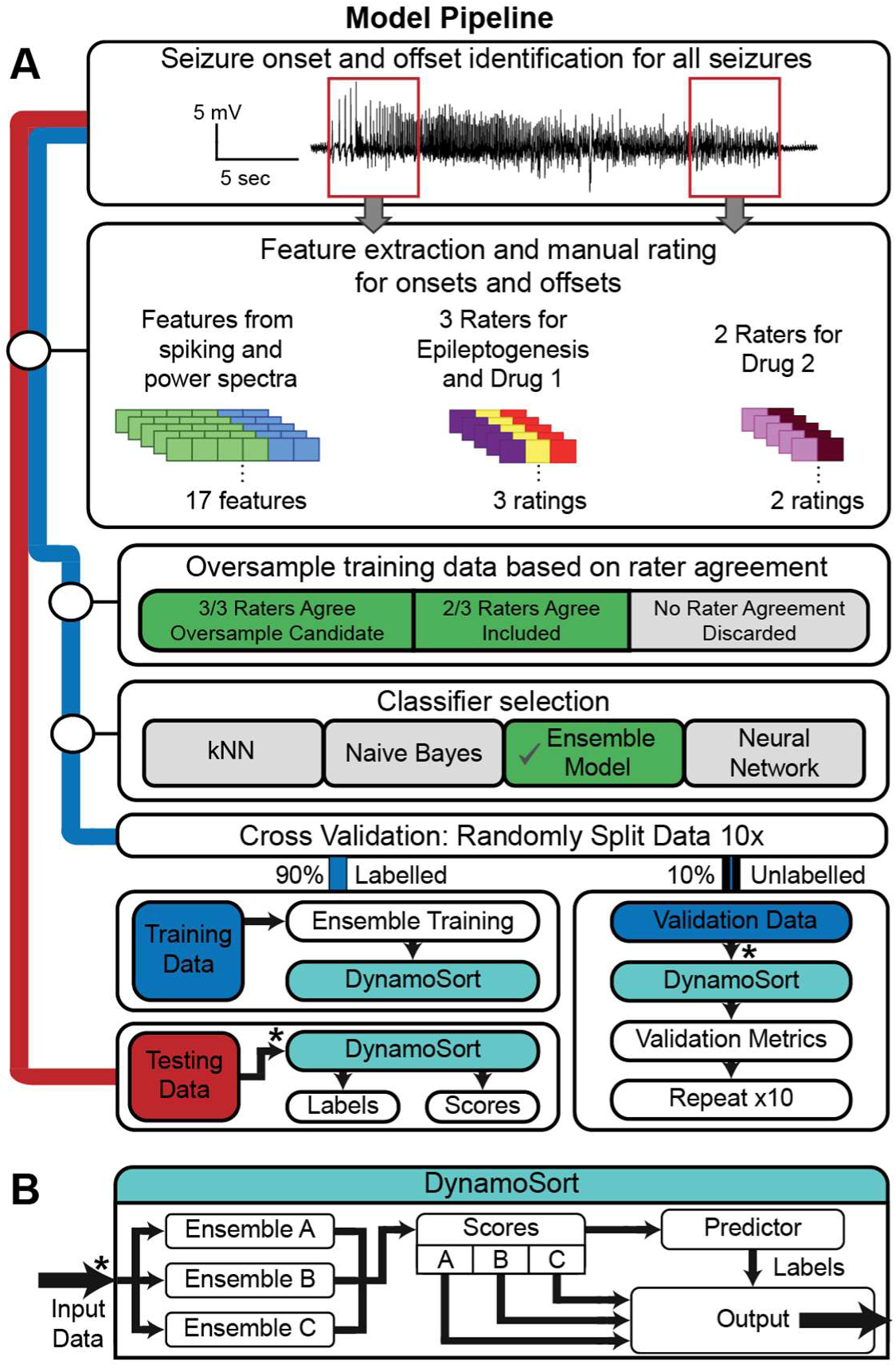
Model Pipeline. **(A)** A flowchart illustrating the processing pipeline for EEG data within the DynamoSort model. Seizure onset and offset windows were segmented for analysis and underwent a series of feature extraction steps. Extracted features were labelled by either Raters A, B, and C, or Raters D and E, depending on the dataset. Training data were used to select the classifier via MATLAB’s Classification Learner application. Once the classifier was selected, the training data were used to train and validate DynamoSort, while testing data were kept separate and exclusively used to evaluate model performance post-training. (**B)** A depiction of the internal operations of the DynamoSort model.

## Results

### Bifurcation type distribution and rater agreement

For the epileptogenesis training dataset, 17% of onsets were SupH, 15% were SNIC, 65% were SN(-DC)/SubH, and 4% were without consensus (Figure 4A). The distribution of the offset data was slightly different, where 13% of offsets were SupH, 22% were SNIC, 62% were FLC, and 3% were without consensus. The SN(-DC) or SubH and FLC bifurcation types, which lack clear patterns in amplitude or frequency, were represented at the highest rates for both onsets and offsets.

**Figure 4.**
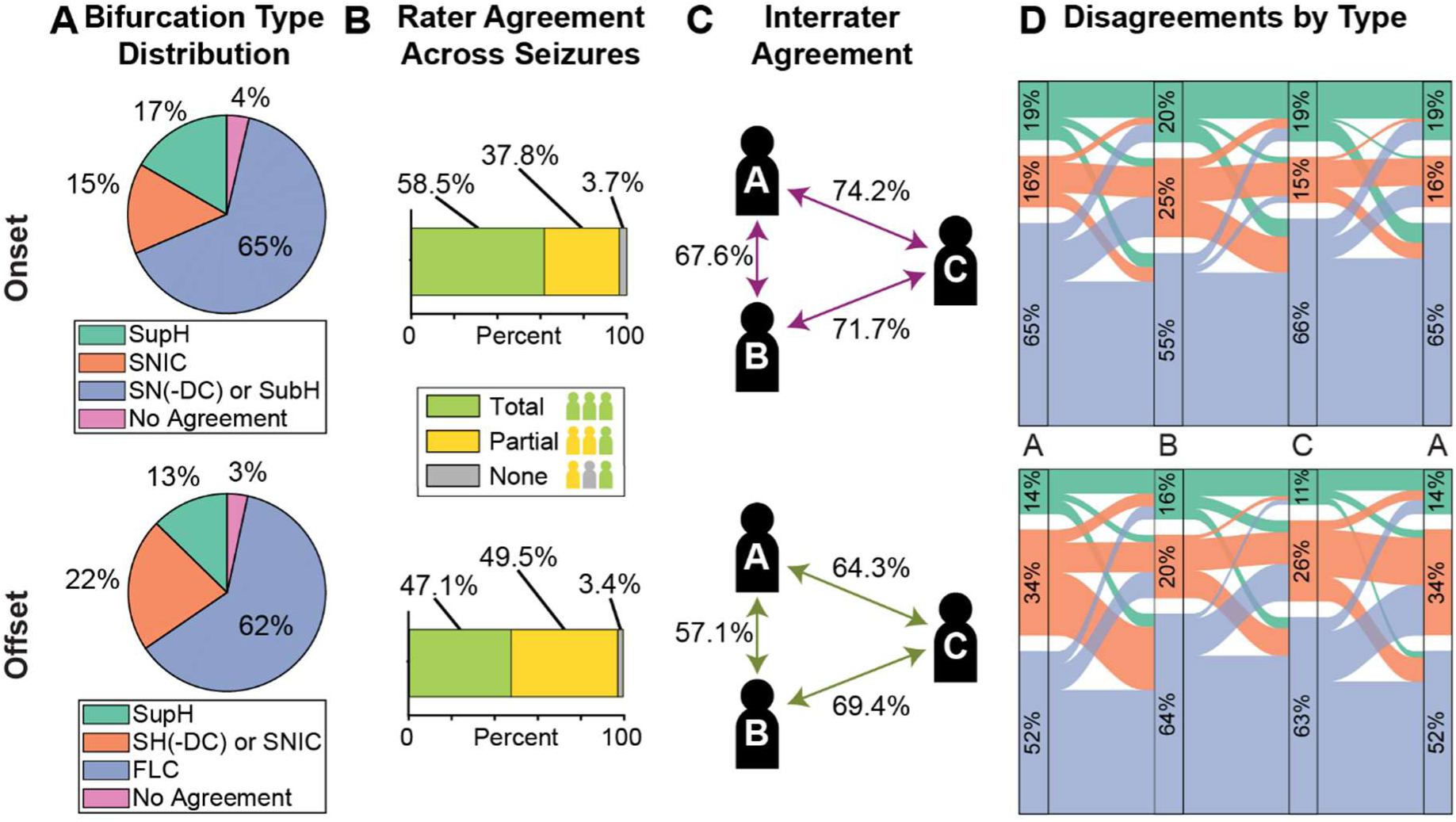
**(A**) Onset and offset distribution of each bifurcation type. Instances where no agreement was made were removed from the training dataset, which was only 3.7% in onset types and 3.4% in offset types. The third category of bifurcation type, characterised by a lack of amplitude or frequency patterning, was most prominent. **(B**) The majority of instances are agreed on by at least two raters for onset types (96.3%) and for offset types (96.6%). **(C)** Agreement between individual raters shown as a percentage of total rating instances, split into onset and offset. Rater C was found to agree with both other raters more often than A and B agreed with each other. There was more disagreement in the offset data (average agreement: 64.2%) than onset data (average agreement: 73.4%). **(D)** Alluvial plot showing each individual rater’s bifurcation type distribution and disagreement trends. Distributions were significantly different (Onset: χ2(4) = 32.68, p < 0.001; Offset: χ2(4) = 41.33, p < 0.001), but effect sizes in both onset and offset were small (Onset: V < 0.1, Offset: V < 0.1).

At least two raters agreed with each other in most cases, with full disagreement under 4% for both onset and offset (Figure 4B). Notably, there was a higher proportion of unanimous consensus for onset bifurcation types (58.5%) compared to offset (47.1%). Interrater agreement was calculated to determine accuracy and consistency across individual raters. Average interrater agreement was higher for onset types (73.4%) than for offset types (64.2%), further illustrating higher uncertainty in classifying offsets (Figure 4C). A chi-square test showed differences in ratings were significant to each rater, but the differences were relatively that the differences in ratings were significant to each rater, but the differences were relatively small (Onset: χ2(4) = 32.68, p < 0.001, V = 0.088; Offset: χ2(4) = 41.33, p < 0.001, V = 0.099). The most frequent disagreements were between frequency-based bifurcations (SNIC and SN(-DC) or SubH) and arbitrary bifurcation types (SN(-DC) or SubH and FLC) in both onset and offset classifications (Figure 4D).

### Model and training dataset validation

We used the ensemble classifier (logitBoost) to develop DynamoSort. This model was the highest performer based on the chosen metrics (Onset mean AUC: 0.81; Offset mean AUC: 0.75) (Figure 5A). The oversampled epileptogenesis dataset was used in the final implementation of the classifier and to gather performance data on the two drug datasets.

**Figure 5.**
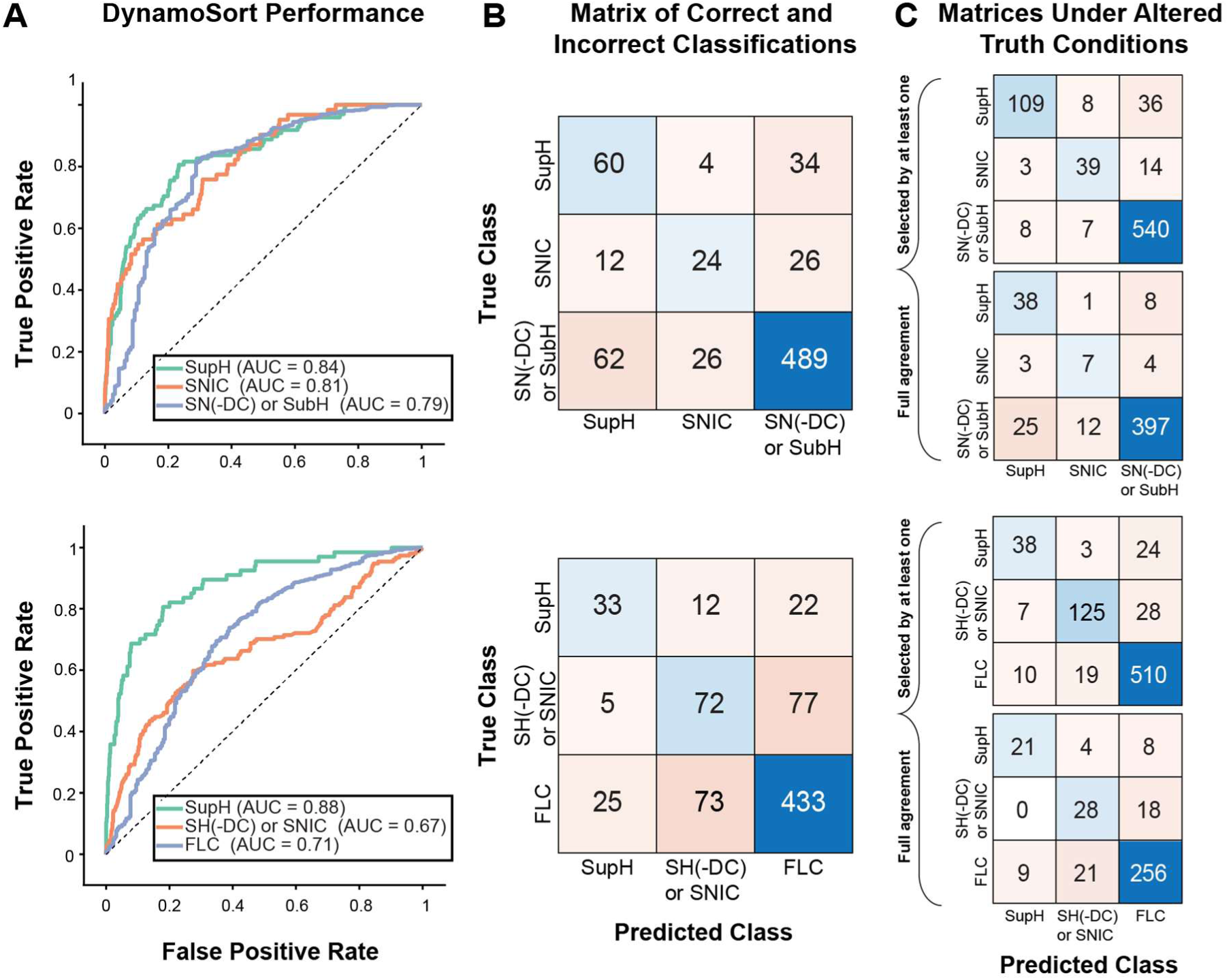
DynamoSort Performance. **(A)** ROC curves with performance (AUC values) listed by bifurcation type for onset and offset. Onset classification (0.79-0.84) performed better on average than offset classification (0.67 – 0.88), but the offset model performs better in classifying SupH bifurcations (0.88 versus 0.84). **(B)** Confusion matrices breakdown the types of misclassifications in the model, such as false positives or false negatives. The confusion matrices visualise performance under the default ground truth conditions, where at least two raters agree on the bifurcation type. True positive rate (TPR) is defined as # of true positives divided by the total number of cases of that type and is shown along the diagonal. For instance, onset SupH had 60 true positives out of 98 total cases, so its TPR would be 0.61. SN(-DC)/SubH and FLC had the highest TPR (0.90 and 0.82) and lowest true negative rates (0.66 and 0.66) for onsets and offsets, respectively. **(C)** In alternative conditions (at least 1 rater classified or full agreement), the TPR greatly improves. The average onset TPR increases from 0.65 under default conditions to 0.79 (at least 1 rater condition) and 0.74 (full agreement condition), and the average offset TPR increases from 0.59 to 0.77 and 0.72 respectively.

### DynamoSort performance

DynamoSort shows high performance on the Drug 1 dataset (mean onset AUC: 0.81, mean offset AUC: 0.75). A confusion matrix was created to visualise the true positive rate (TPR: the algorithm correctly classifies the bifurcation type) and true negative rate (TNR: the algorithm correctly classifies the type as the negative type) for the Drug 1 test dataset (Figure 5B). With three bifurcation type possibilities—for both onset and offset independently—a TPR and TNR of 0.33 would signify that the model performs no better than chance in classification and 1 would signify perfect classification. The initial average onset TPR was 0.65, and the initial average offset TPR was 0.59 (Supplemental Table 2). These results, while better than chance, did not initially appear to be strong enough for fully automatic classification. However, just as human raters tend to disagree with each other (57.0%—78.8% of the time in our datasets), the “ground truth” as previously defined may not fully illustrate the model’s capabilities.

To better quantify model performance with the understanding that even human raters can disagree with each other, we explored “alternative truth” conditions (Figure 5C). First, we evaluated the algorithm’s performance using a more liberal truth definition, where an onset or offset classification was valid if at least one human rater identified it as such (e.g. when two raters labelled an onset as SupH, and the third labelled it as SNIC, both SupH and SNIC would be acceptable “true” labels). Under these conditions, DynamoSort’s performance expectedly increases (Supplemental Table 2). The average onset TPR rises from 0.65 under default conditions to 0.79. Importantly, these gains are not only from one bifurcation type. Offset average TPR also rose under the new conditions, from 0.59 to 0.77 (minimum from 0.47 to 0.59).

The other alternative “truth” condition was to evaluate performance for the onset and offset classification in cases when all three of the raters agreed. These unanimous classifications were determined to be strong examples of each bifurcation type. As in the other alternative truth condition, performance increased for all types and on average. The average onset TPR rose from 0.65 to 0.74, and the average offset TPR rose from 0.59 to 0.72.

### DynamoSort/human agreement and bifurcation likelihood scoring

Comparing interrater agreement between raters A, B, and C in the epileptogenesis dataset and agreement between raters D, E and DynamoSort (i.e., rater F) in the Drug 2 dataset; there are clear parallels (Figure 4C and Figure 6A). A Fisher’s exact test showed there was no statistically significant difference in human-human agreement of D and E (ABC-DE: p=0.21, 95% CI=0.92-1.46) or in human-machine agreement (ABC-DF: p=0.59, 95% CI=0.0.74-1.19; ABC-EF: p=0.34, 95% CI=0.0.89-1.41). For offsets, there was a significant difference between the human-machine agreement between rater E and DynamoSort than average human-human agreement across ABC raters (ABC-EF: p<0.001, 95% CI=1.18-1.82). However, there were also cases of significantly different levels of agreement in *just* human raters in the training dataset, such as the agreement between A and C versus the agreement B and C in the offset (AC-BC: p<0.00001, 95% CI=0.47-0.73).

**Figure 6.**
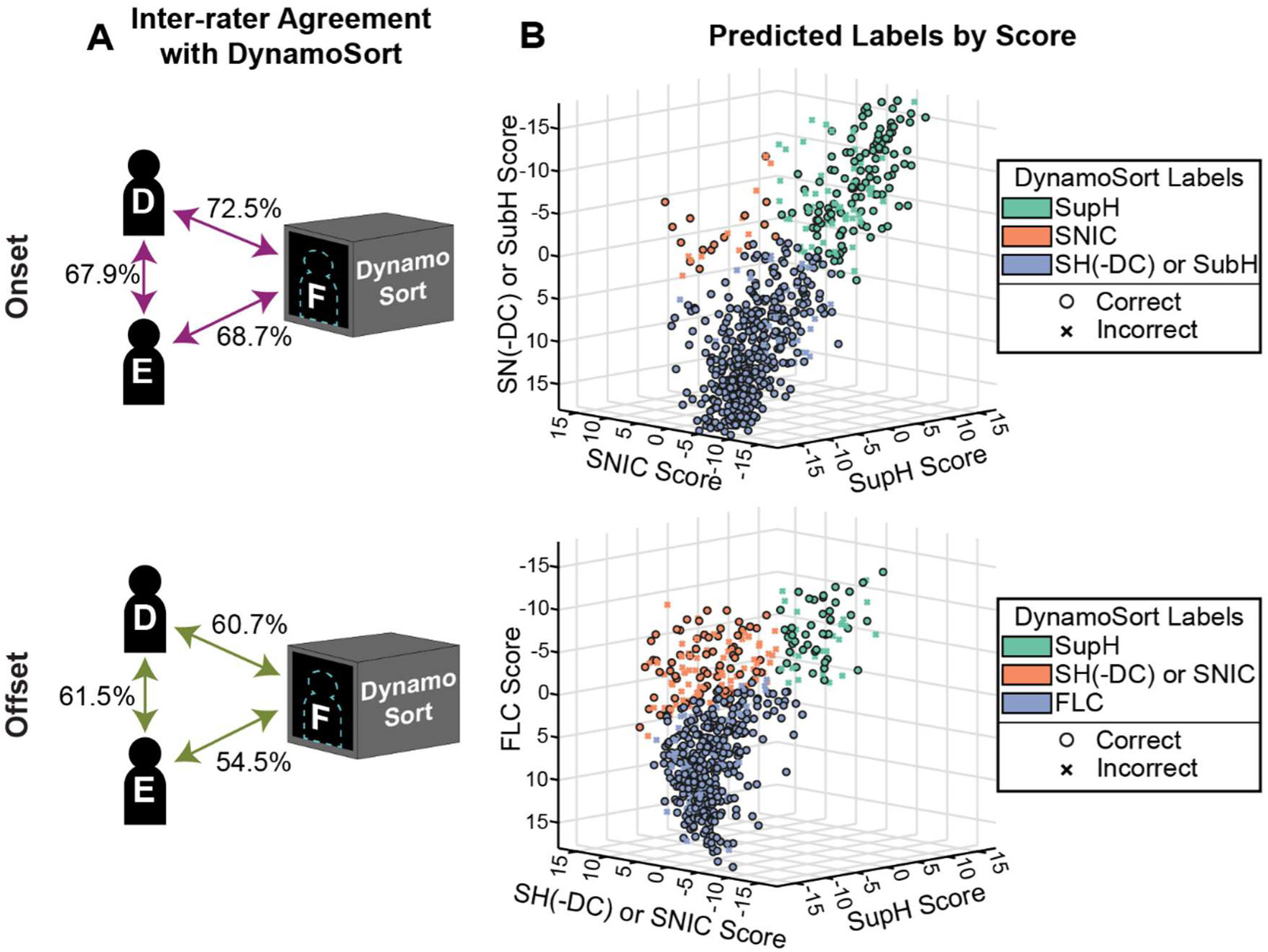
**(A)** In the second test dataset, agreement between two different human raters, D and E, was 67.9% and 61.5% for onset and offset types, respectively. Agreement between DynamoSort and human raters D and E averaged 70.6% and 57.6% for onset and offset types, respectively. The percentages exhibited parallel trends with the level of human disagreements and were not significantly different for onsets (p = 0.59, Fisher’s exact test), and while the disagreements in offsets were significantly greater (p<0.001, Fisher’s exact test), there were instances of human-human disagreement for offset classification that were also significantly different (p<0.00001, Fisher’s exact test). **(B)** 3D scatterplot showing the DynamoSort bifurcation likelihood score distribution, coloured by subtype and annotated by agreement with human classification. Clear clusters emerge showing numerical separation of bifurcation types.

A key feature of DynamoSort is its ability to assign probability scores for each bifurcation type independently, providing a quantitative analysis of onset and offset shape. Figure 6B shows DynamoSort scores for each seizure in the Drug 2 dataset, along with how its classification aligns with rater labels. Because the Drug 2 dataset had only 2 raters, a “true” case was defined as one rater labelling an instance as that type (e.g. if rater D chose SupH and rater E chose FLC, the DynamoSort’s classification was considered correct if either type was chosen), whereas a “false” case meant neither rater chose that specific bifurcation type. Across all bifurcation types, the probabilistic scores were significantly higher for true classifications than false cases (Mann-Whitney U test, p<0.005). The largest difference was observed in the SN(-DC)/SubH onset type (mean true score: 5.31; mean false score: -5.22).

These scores directly relate to the feature values extracted from the EEG (Figure 7). To quantify this relationship, we performed pair-wise Mann-Whitney U tests between each bifurcation type and feature. Across the 17 features, each significantly contributed to distinguishing dynamotypes. Overall, DynamoSort mirrored human classification, with amplitude features being more crucial for classifying SupH onset and offset subtypes, while frequency-related features were more influential for classifying SNIC onsets and SH(-DC)/SNIC offsets.

**Figure 7.**
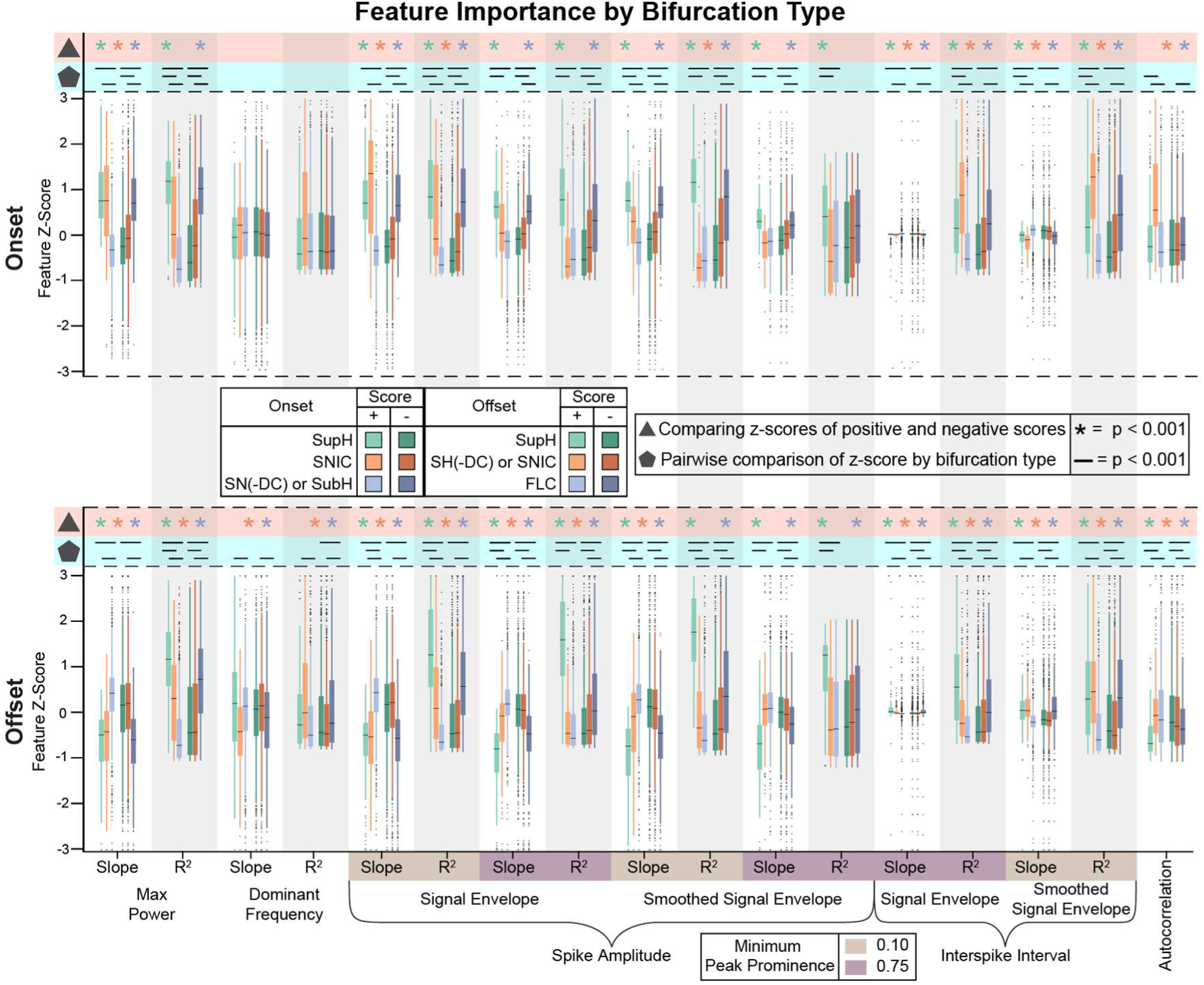
DynamoSort scores correspond to critical features for each bifurcation type. The DynamoSort bifurcation likelihood score for each subtype can range from -20 to 20, with more positive scores indicating the onset/offset exhibits more features characteristic of that subtype. We evaluated significant features across positive scores (score > 0→ +) and negative scores (score ≤0→ -) for each bifurcation type for seizure onsets and offsets. Features exhibited expected trends, with certain features contributing more prominently to specific bifurcation type selections. For example, the 0.1 Smoothed Amplitude slope feature was significantly higher in SupH onsets than SNIC (p < 0.0005) and SN(-DC)/SubH (p < 0.0005). Positive and negative scores were compared for each feature and by type using a Mann-Whitney U test, corrected for multiple comparisons, which was also used to compare bifurcation types directly to each other.

## Discussion

In the present study, we developed DynamoSort to automatically classify seizures based on bifurcation types. We successfully trained, validated, and tested this model, which performs comparably to human raters. Using 17 features, the model was able to score seizure onsets and offsets according to their dynamics.

### Distribution of bifurcation types

For EEG seizure data where the DC offset was filtered out, as seen in the three datasets used in this study, a maximum of 3 onset and 3 offset bifurcation types could be distinguished. All types were present in our analysis, with distributions similar to what has been reported in humans (Saggio et al. 2020). With our training dataset, SupH : SNIC : SN(-DC)/SubH types were represented in 17% : 15% : 65% of the seizures and SupH : SH(-DC)/SNIC : FLC were represented in 13% : 22% : 62% of the seizures. In seizures recorded intracranially in humans, Saggio et al., observed that SupH : SNIC : SN(-DC)/SubH types were represented in 26% : 6% : 67% of the seizures and SupH : SH(-DC)/SNIC : FLC were represented in 3% : 41% : 54% of the seizures. The arbitrary bifurcation types (SN(-DC)/SubH and FLC) were the most common bifurcation types across onsets and offsets for both mouse and human seizures. Interestingly, there was a difference in the representation of the frequency-based bifurcation types (onset SNIC and offset SH(-DC)/SNIC types). Saggio et al. found that SH(-DC)/SNIC types were nearly twice as common in offsets in human recordings than we found in mice. While rater bias may have played a role in these differences, the use of multiple raters helped mitigate this possibility. However, there may be a fundamental difference between onset and offset type distributions in a human cohort containing multiple epilepsy aetiologies compared to IAK mice, an induced model of mesial temporal lobe epilepsy.

Differences in distributions may be related to the mechanisms of seizure initiation and termination. In a preprint, experiments by Crisp et al., *in vitro* brain slices using a high magnesium, no potassium excitatory media observed the ratio of SupH : SNIC : SN(-DC)/SubH onset bifurcation types to be in 12% : 63% : 24%, with a notable prominence of the frequency-based SNIC onset type compared to the prominent SN(-DC)/SubH type in the *in vivo* data collected in our manuscript and Saggio et al.,2020 (Crisp, Parent, et al. 2020, Saggio et al. 2020). However, *in vitro*, the ratio of SupH : SNIC : SN(-DC)/SubH types changed to 35% : 12% : 54% in the presence of anti-seizure drug GABA and to 43% : 43%: 13% in the presence of anti-seizure drug carbamazepine. The differential changes to onset types in the presence of these drugs might be explained by their distinct mechanisms of action: GABA is a chloride ion channel blocker (Davies 1995). This suggests that the presence of specific bifurcation types may be indicative of different brain states, each leading to seizure onset via different pathways.

Finally, Guendelman et al. 2025 conducted seizure dynamotype classification using non-invasive recordings to test applicability to surface EEG. They used an automated random forest classifier and Bayesian estimator to classify dynamotypes as defined by criteria in Saggio et al, 2020. Of the 1,177 seizures analysed from the EPILEPSIAE database (Ihle et al. 2012), 49.5% exhibited clear onset bifurcations and 40.3% exhibited clear offset bifurcations. For onset bifurcations, the SupH : SNIC : SN(-DC)/SubH distributions were 6.7% : 4.6% : and 88.7%, and offset distributions for SupH : SH(-DC)/SNIC : FLC were 0.6% : 28.7% : 70.7%. These non-invasive recordings show a clear predominance to the third bifurcation type (SN(-DC)/SubH and FLC), which may indicate differences in differentiating bifurcation types between scalp versus intracranial recordings, possibly due to filtering of high frequency oscillations through the skull (Zelmann et al. 2014), with the latter possibly capturing more amplitude- and frequency-dependant bifurcation types.

### Interrater Agreement

The dynamotype classification method is descriptive and based on an oscillator model rather than distinct biological mechanisms, so no ground truth exists for classification.

Disagreement between raters is expected, and in our analysis, onset bifurcations were 58.5% unanimous, 37.8% with 2/3 rater agreement, and 3.7% had no consensus, while offset bifurcations were more challenging (47.1% unanimous, 49.5% 2/3 agreement, 3.4% no consensus).

Levels of interrater agreement for onset bifurcations in Crisp et al., 2020 were 62% unanimous, 36% 2/3 agreement, and 2% no consensus. Consistency across raters using the taxonomy defined in Saggio et al, 2020 were similar, with of no consensus ranging from 1— 3%. Interestingly, in an experiment in Saggio et al., 2020, where simulated data was used to test rater agreement—where a ground truth does exist—raters were 80% unanimous, 18.3% 2/3 agreement, and 1.7% no consensus for onset types and 78.75% unanimous, 21.25% 2/3 agreement, 0% no consensus for offset types. Notably, disagreement across raters in biological data may not indicate errors but rather reflect raters observing and prioritising features of multiple bifurcation types during their ratings. DynamoSort performed comparably to a human rater but goes a step further by providing a score on a continuum to represent these potential grey areas in classification.

### DynamoSort performance

The algorithm demonstrates robust performance in both onset and offset classification, with a mean AUC exceeding 0.7 in both cases (0.81 for onsets and 0.75 for offsets, Figure 5A)— generally considered the standard threshold for this type of classification (Nahm 2022). This level of performance indicates the model is suitable for supervised sorting and can serve as a “tiebreaker” between raters. Moreover, when the classification was considered correct if at least 1 rater chose that type, or in full agreement cases, performance of the algorithm was greatly improved, as demonstrated in Figures 5B and 5C, where the true positive rate increased up to 14% for onsets and up to 18% for offsets. As stated, onset and offset classifiers remained separate throughout all training, validating, and testing. The differences between the two classifiers were in specific hyperparameter optimizations and users can combine the separate onset and offset types to obtain the full dynamotype.

### Seizure complexity

A known limitation in dynamotype classification is applying low-dimensional mathematical models to systems as complex as the brain. We have shown that while at least two raters will usually agree with each other, disagreement on specific bifurcation type is not uncommon. Some disagreements can be explained by human error, but others may be due to multiple bifurcation types being present at once. While some of these bifurcations cannot feasibly co-occur in mathematics, the co-occurrence or “jumping” between bifurcation types has been noted in the literature (Saggio et al. 2020; Crisp, Cheung, et al. 2020). We observed the same in our datasets, with DynamoSort assigning positive likelihood scores to 2 out of 3 bifurcation types for some seizures. It has also been found that the dominant bifurcations within individuals can evolve over time in rodent models (Crisp, Cheung, et al. 2020). The use of bifurcation likeliness scores will enable the analysis of changing bifurcations across both long timescales (throughout epileptogenesis or an individual’s life) and short timescales (e.g., multiple bifurcations within a single seizure). Rather than simply observing a sudden shift from one dominant bifurcation to another, we can now quantify the progression of these transitions, which could be especially useful in biomarker analysis.

Dynamotypes have shown promise as a biomarker in both the treatment and research of epilepsy. It has been demonstrated that bifurcation types influence the effectiveness of seizure aborting stimulation in computational models (Szuromi et al. 2023), with the relationships between the limit cycle and resting states affecting the stimulation parameters required to terminate an epileptic burst. The same dynamics have been employed in network analysis to help determine the epileptogenic zone in focal epilepsy patients (Runfola et al. 2023), and have been incorporated into virtual brain models for the same zone-identification purpose (Jirsa et al. 2023). Finding the epileptogenic zone, where seizures originate in focal epilepsy, is critical for successful resection surgery, and dynamical mechanisms have proven valuable in enhancing this process. As there is clearly interest in furthering seizure dynamics research under the dynamotype taxonomy, we developed DynamoSort as a means to quantify these seizure dynamics without the high cost of manual labelling.

### Future Directions

Although the dynamic networks themselves were not within the scope of this work, prior defined seizure bifurcation types (Saggio et al., 2020) were observed in the IAK mouse model. While the dynamotype taxonomy was created with humans in mind, its application in mouse models allows more flexibility in their study as a biomarker.

With the recent increase in epilepsy research focused on network change, biomarkers such as EEG signal onset and offset characteristics—easily categorized by the dynamotype taxonomy—are critically important. For example, studies have shown altered oscillations during and around sleep in the epileptic brain, and that seizures often occur according to circadian and multidien rhythms (Bernard et al. 2023, Bernard 2021, Baud et al. 2018, Karoly et al. 2021, Anderson et al. 2024, Charlebois et al. 2024). However, the way that these oscillations and rhythms change over time remains unclear due to the limited ability to continuously monitor EEG over long time scales. Beyond preliminary *in vitro* studies by Crisp, Cheung, et al., 2020, there is also emerging evidence that bifurcation types change based on brain state, such as during sleep. Using non-invasive data, Guendelman et al., 2025, found that dynamotypes changed based on sleep state, where SNIC and SupH onset types had greater prevalence in seizures during wake and REM sleep. Additionally, SupH was observed more during focal impaired awareness seizures (6.2%) compared to focal aware seizures (1.8%) or focal-to-bilateral-tonic-clonic seizures (0%).

A better understanding of brain states is becoming an important topic in epilepsy research, especially in the context of state-dependent neurostimulation in epilepsy. Experiments have been conducted *in silico* to optimize ictal abortive stimulation (Salam et al. 2016), in rodents (Zhang et al. 2023), and human studies show outcomes of patients with closed-loop neurostimulation devices may depend on the brain state in which they received stimulation (Anderson et al. 2024). Certain dynamotypes may be more responsive to ASMs or stimulation approaches. Szuromi et al, 2023 found seizures with a baseline shift (DC offset) were more likely to be aborted with stimulation in computational simulations. As this field evolves, DynamoSort offers an easy-to-use machine learning algorithm capable of generating both discrete labels and continuous scores, facilitating meaningful analysis of such biomarkers in future work.

### Limitations

In the creation of DynamoSort, and many models like it, a major limiting factor is dataset size. Training researchers to label large datasets is time consuming and expensive, which was the original rationale behind building DynamoSort. Lack of data and imbalanced types means that there is a likelihood that some of the false positives and false negatives are the result of overtraining. We accounted for this by oversampling strong instances of the bifurcation types that were expressed less frequently, such as SNIC and SupH, which we believe helps protect against some imbalance. While this could be seen as a limitation, the algorithm extracts features from spiking activity and time-frequency spectrograms in a manner that is agnostic to the data’s original source. Since dynamotypes are classified based on changes in signal amplitude and frequency, DynamoSort is expected to perform effectively on other data sources as well.

Another major limitation is the lack of an objective ground truth of classification labels in our dataset. When training a machine learning classifier, success depends largely on the training data it is given. While complete disagreement was relatively low for both onsets and offsets (3.4%–3.7% the proportion of instances where at least one rater disagreed was high (37.8%–49.5%). Nearly 50% of cases having at least one dissenter is evidence of how difficult this task can be even for trained human raters. Despite this, the presence of disagreement provided valuable insights into the algorithm performance and was comparable to human performance. In some cases, disagreement may arise because seizure onset and offset patterns exhibit traits of multiple bifurcation types. This motivated the development of DynamoSort’s ability to produce a likelihood score, acknowledging that black-and-white classification may not be feasible using EEG seizure data. As an openly available model, however, DynamoSort can be continuously updated with new training data to improve its accuracy and adaptability over time. It can also be directly compared with new ratings and tuned to label groups according to needs.

## Author Contributions

JW contributed conceptualization, methodology, software, formal analysis, investigation, writing – original draft, visualization. AZ contributed methodology, validation, data curation, writing – review and editing. ML contributed data curation. SS contributed data curation. KJ contributed data curation. JMR contributed methodology and writing – review and editing. PJW contributed conceptualization and resources. KSW contributed conceptualization, resources, writing – review and editing, supervision, and funding acquisition. DNA contributed conceptualization, methodology, software, formal analysis, resources, data curation, writing – review and editing, visualization, supervision, funding acquisition.

## Data Availability

The DynamoSort code and model will be available upon publication. Mouse EEG data will be available upon request.

## Ethics Statement

Data used to train and test DynamoSort from West et al., *Exp. Neurol.*, 2021 were acquired with approval from the University of Utah’s Institutional Animal Care and Use Committee.

## References

Anderson, DN et al. 2024, “Closed-loop stimulation in periods with less epileptiform activity drives improved epilepsy outcomes,” Brain, vol. 147, no. 2, pp. 521–531.

Baud, MO et al. 2018, “Multi-day rhythms modulate seizure risk in epilepsy,” Nature Communications, vol. 9, no. 1, p. 88.

Bernard, C 2021, “Circadian/multidien Molecular Oscillations and Rhythmicity of Epilepsy (MORE),” Epilepsia, vol. 62, no. S1.

Bernard, C et al. 2023, “Sleep, oscillations, and epilepsy,” Epilepsia, vol. 64, no. S3, pp. S3–S12, viewed 29 May 2024, <https://onlinelibrary.wiley.com/doi/full/10.1111/epi.17664>.

Bragin, A et al. 2007, “Analysis of Initial Slow Waves (ISWs) at the Seizure Onset in Patients with Drug Resistant Temporal Lobe Epilepsy,” Epilepsia, vol. 48, no. 10, pp. 1883–1894.

Charlebois, CM et al. 2024, “Circadian changes in aperiodic activity are correlated with seizure reduction in patients with mesial temporal lobe epilepsy treated with responsive neurostimulation,” Epilepsia, vol. 65, no. 5, pp. 1360–1373.

Crisp, DN, Cheung, W, et al. 2020, “Quantifying epileptogenesis in rats with spontaneous and responsive brain state dynamics,” Brain Communications, vol. 2, no. 1.

Crisp, DN, Parent, R, et al. 2020, “Carbamazepine and GABA have distinct effects on seizure onset dynamics in mouse brain slices.”

Davies, JA 1995, “Mechanisms of action of antiepileptic drugs,” Seizure, vol. 4, no. 4, pp. 267–271.

Elahian, B et al. 2018, “Low-voltage fast seizures in humans begin with increased interneuron firing,” Annals of Neurology, vol. 84, no. 4, pp. 588–600.

Guendelman, M, Vekslar, R, & Shriki, O 2025, “A New Perspective in Epileptic Seizure Classification: Applying the Taxonomy of Seizure Dynamotypes to Noninvasive EEG and Examining Dynamical Changes across Sleep Stages,” eneuro, vol. 12, no. 1, p. ENEURO.0157-24.2024.

Ihle, M et al. 2012, “EPILEPSIAE – A European epilepsy database,” Computer Methods and Programs in Biomedicine, vol. 106, no. 3, pp. 127–138.

Jin Huang & Ling, CX 2005, “Using AUC and accuracy in evaluating learning algorithms,” IEEE Transactions on Knowledge and Data Engineering, vol. 17, no. 3, pp. 299–310.

Jirsa, V et al. 2023, “Personalised virtual brain models in epilepsy,” The Lancet Neurology, vol. 22, no. 5, pp. 443–454.

Jirsa, VK et al. 2014, “On the nature of seizure dynamics,” Brain, vol. 137, no. 8, pp. 2210–2230.

Karoly, PJ et al. 2021, “Cycles in epilepsy,” Nature Reviews Neurology, vol. 17, no. 5, pp. 267–284.

Maheshwari, A 2020, “Rodent EEG: Expanding the Spectrum of Analysis,” Epilepsy Currents, vol. 20, no. 3, pp. 149–153.

Milior, G et al. 2023, “Animal models and human tissue compared to better understand and treat the epilepsies,” Epilepsia, vol. 64, no. 5, pp. 1175–1189.

Moshé, SL et al. 2015, “Epilepsy: new advances,” The Lancet, vol. 385, no. 9971, pp. 884–898.

Nahm, FS 2022, “Receiver operating characteristic curve: overview and practical use for clinicians,” Korean Journal of Anesthesiology, vol. 75, no. 1, pp. 25–36.

Noachtar, S & Rémi, J 2009, “The role of EEG in epilepsy: A critical review.”

Runfola, C et al. 2023, “In pursuit of the epileptogenic zone in focal epilepsy:a dynamical network biomarker approach,” Communications in Nonlinear Science and Numerical Simulation, vol. 117, p. 106973, <https://www.sciencedirect.com/science/article/pii/S1007570422004609>.

Rusina, E, Bernard, C, & Williamson, A 2021, “The Kainic Acid Models of Temporal Lobe Epilepsy,” eneuro, vol. 8, no. 2, p. ENEURO.0337-20.2021.

Saggio, ML et al. 2017, “Fast–Slow Bursters in the Unfolding of a High Codimension Singularity and the Ultra-slow Transitions of Classes,” The Journal of Mathematical Neuroscience, vol. 7, no. 1, p. 7.

Saggio, ML et al. 2020, “A taxonomy of seizure dynamotypes,” eLife, vol. 9, p. e55632, <10.7554/eLife.55632>.

Salam, MT, Perez Velazquez, JL, & Genov, R 2016, “Seizure Suppression Efficacy of Closed-Loop Versus Open-Loop Deep Brain Stimulation in a Rodent Model of Epilepsy,” IEEE Transactions on Neural Systems and Rehabilitation Engineering, vol. 24, no. 6, pp. 710–719.

Staba, RJ, Stead, M, & Worrell, GA 2014, “Electrophysiological Biomarkers of Epilepsy,” Neurotherapeutics, vol. 11, no. 2, pp. 334–346, <https://www.sciencedirect.com/science/article/pii/S1878747923009029>.

Szuromi, MP, Jirsa, VK, & Stacey, WC 2023, “Optimization of ictal aborting stimulation using the dynamotype taxonomy,” Journal of Computational Neuroscience, vol. 51, no. 4, pp. 445–462.

The MathWorks Inc. 2023, “MATLAB.”

West, PJ et al. 2022, “Spontaneous recurrent seizures in an intra-amygdala kainate microinjection model of temporal lobe epilepsy are differentially sensitive to antiseizure drugs,” Experimental Neurology, vol. 349, p. 113954.

Zelmann, R et al. 2014, “Scalp EEG is not a Blur: It Can See High Frequency Oscillations Although Their Generators are Small,” Brain Topography, vol. 27, no. 5, pp. 683–704.

Zhang, F et al. 2023, “Efficacy of different strategies of responsive neurostimulation on seizure control and their association with acute neurophysiological effects in rats,” Epilepsy & Behavior, vol. 143, p. 109212.

