## Supplemental Tables 1 & 2 for "DynamoSort: Using machine learning approaches for the automatic classification of seizure dynamotypes"

| Dataset | Seizures | Animals | Raters | Training | Validation | Testing |
|---|---|---|---|---|---|---|
| Epileptogenesis | 709 | 22 | A, B, C | X | X | |
| Drug 1 | 765 | 18 | A, B, C | | | X |
| Drug 2 | 638 | 14 | D, E | | | X |

*Supplemental Table 1.* ***Distribution of data sets across training, validation, and testing phases.*** *The epileptogenesis and Drug 1 datasets all underwent the same manual labelling process, by raters A, B, and C. The Drug 2 dataset underwent an identical labelling process by raters D and E. The epileptogenesis seizure set (709 seizures) was used in training and cross-validation, while Drug 1 (765 seizures) and Drug 2 (638 seizures) datasets were used to test efficacy of the model.*

| Truth conditions | | Default majority vote conditions | | Selected by at least one rater | | Full agreement conditions | |
|---|---|---|---|---|---|---|---|
| | | TPR | TNR | TPR | TNR | TPR | TNR |
| Onset | SupH | 0.61 | 0.90 | 0.71 | 0.98 | 0.81 | 0.94 |
| | SNIC | 0.44 | 0.96 | 0.70 | 0.98 | 0.50 | 0.95 |
| | SN(-DC)/SubH | 0.90 | 0.66 | 0.97 | 0.77 | 0.91 | 0.82 |
| | Mean | 0.65 | 0.84 | 0.79 | 0.91 | 0.74 | 0.90 |
| Offset | SupH | 0.49 | 0.96 | 0.59 | 0.98 | 0.64 | 0.98 |
| | SH(-DC)/SNIC | 0.47 | 0.86 | 0.78 | 0.98 | 0.62 | 0.92 |
| | FLC | 0.82 | 0.66 | 0.95 | 0.86 | 0.89 | 0.90 |
| | Mean | 0.59 | 0.83 | 0.77 | 0.94 | 0.72 | 0.93 |

*Supplemental Table 2.* ***True and false positive rates for each bifurcation type in each ground truth condition.*** *The true positive rate (TRP) and true negative rate (TNR) were used as metrics to analyse model performance. The TPR and TNR for each bifurcation type were shown under three "truth" conditions, along with means for each. TPR and TNR both increase under alternative-truth conditions, with a maximum improvement of +0.31 (offset SH(-DC)/SNIC from 0.47 TPR to 0.78 TPR). These alternative-truth conditions account for disagreements found in human rater datasets.*
